# Persistent CCL3 expression sustains neutrophil recruitment and contributes to delayed wound healing in advanced age

**DOI:** 10.64898/2026.05.28.727592

**Authors:** Holly R Rocliffe, Yonlada Nawilaijaroen, Velizar Shinkov, Antonella Pellicoro, Sarah R Walmsley, Jenna L Cash

## Abstract

Delayed wound healing in advanced age is associated with prolonged inflammation, but the mechanisms that impair inflammatory resolution in this context remain poorly defined. Neutrophils are essential for early repair, but their persistence is predicted to disrupt resolution and delay healing. Here we show that aged mice exhibit impaired wound closure accompanied by sustained neutrophil infiltration from days 7–14 post-injury, long after neutrophils have resolved in young wounds. In aged skin, macrophage abundance and neutrophil–macrophage interactions were reduced, consistent with reduced macrophage-mediated clearance. Instead, neutrophils preferentially localised to perivascular regions, consistent with ongoing recruitment. Transcriptomic profiling demonstrated that young wounds transitioned from inflammatory to reparative programs, whereas aged wounds retained inflammatory signatures, including enrichment for neutrophil chemotaxis and LPS-response pathways. Notably, *Ccl3, Cxcl2* and *Cxcl3* remained elevated in aged wounds. Analysis of human chronic ulcers identified macrophages as a major source of these transcripts, and aged mouse wounds possessed more CCL3+ macrophages than in young. Targeted blockade of CCL3 beginning at day 4 post-injury reduced neutrophil accumulation, accelerated wound closure, and rescued key features of age-impaired healing, when administered after the initial inflammatory phase.

Together, these findings identify persistent CCL3 expression as a contributor to sustained neutrophil recruitment and impaired repair in advanced age. Therapeutically targeting this pathway offers a strategy to restore timely resolution and improve healing outcomes.

## INTRODUCTION

Cutaneous wound healing is a highly coordinated process involving sequential yet overlapping phases of haemostasis, inflammation, proliferation and remodelling (1,2). Successful repair depends not only on the initiation of these processes, but on their timely resolution; disruption of the magnitude, duration or spatial organisation of repair-associated responses can lead to delayed healing or chronic non-healing wounds (3).

Inflammation is an essential early component of wound repair, providing antimicrobial defence and enabling debris clearance. Neutrophils are among the first immune cells recruited to the wound bed, responding to chemotactic signals including CCL2, CCL3, CXCL1 and CXCL8, and mediate host defence through phagocytosis, protease release and production of reactive oxygen species (4,5). Experimental studies demonstrate that neutrophils are required during the early inflammatory phase, as depletion prior to injury impairs healing, and impaired early neutrophil trafficking contributes to defective repair in diabetic wounds, whereas restoration of early recruitment improves healing outcomes (6,7). However, neutrophil activity must be tightly regulated as excessive or prolonged accumulation promotes tissue damage, impairs re-epithelialisation and delays progression to later stages of healing (8,9). Current understanding suggests that effective repair depends on the timing of neutrophil recruitment rather than simply the magnitude of their infiltration.

Advanced age is associated with delayed wound repair and an increased risk of chronic, non-healing ulcers, reflecting disruption of multiple components of the healing cascade, including keratinocyte migration, angiogenic and immune regulation (10–12). A consistent feature of aged wounds is a prolonged inflammatory phase, characterised in part by persistence of neutrophils within the wound bed. Recent single-cell transcriptomic analyses demonstrate persistent neutrophil populations and sustained pro-inflammatory macrophage programmes in aged wounds, indicating a defect in the transition from inflammatory to reparative phases rather than simple exaggeration of early inflammation (13). However, the mechanisms that sustain neutrophil persistence in aged wounds, and whether these can be therapeutically targeted to promote resolution and improve repair, remain poorly defined.

Chemokines are central regulators of leukocyte recruitment and retention within injured tissues, yet their contribution to age-associated defects in inflammatory resolution has not been systematically explored. Here, we show that advanced age is associated with delayed progression through key stages of wound repair, accompanied by persistent neutrophil accumulation and altered immune cell organisation within the wound bed. Aged wounds retained inflammatory transcriptional programmes, including elevated expression of neutrophil-attracting chemokines such as CCL3. We further demonstrate that transient CCL3 blockade after the early inflammatory response reduces late neutrophil accumulation and improves wound healing outcomes. Together, these findings identify sustained chemokine expression as a contributor to impaired inflammatory resolution during wound repair in advanced age.

## RESULTS

### Aged skin exhibits delays throughout wound healing

Skin is composed of a stratified epidermis, dermis and hypodermis, which in mice is underlain by the panniculus carnosus (muscle) layer. Our 4 mm dorsal excisional wounding model involves complete removal of epidermis, dermis, hypodermis and panniculus carnosus using a punch biopsy tool to create 4 uniform wounds (Fig.1A). Skin wounds in young (8 week old) mice consistently healed faster than their aged (18 month) counterparts, with the most noticeable differences in wound area observed at 3 and 7 days post-wounding (dpw; Fig. 1B-E). At 3 dpw wound area had reduced to 36% of original area in young mice, but 52% in aged mice, whilst at 7 dpw wound area in young mice had further reduced to only 8.3% in comparison to 23.6% in the aged counterparts (Fig.1B-C). One of the first events in wound healing is scab formation due to platelet activation to form a fibrin plug. This temporarily reinstates barrier function at the wound site, allowing re-epithelialisation to occur under the scab, which is then lost once the permanent epidermal barrier has reformed. In aged animals, scab formation and loss were delayed. Scab formation was complete within 24 hours of wounding in young animals, but in aged animals, only 25% of wounds exhibited a scab at this time point (Fig.1D). Conversely, at 10 dpw, the majority of wounds in young animals had lost their scabs, indicating completion of wound re-epithelialisation (scab presence 9% young, 67% aged mouse wounds at 10 dpw; Fig 1D). Consistent with delayed re-epithelialisation, histological analysis demonstrated a significantly larger epithelial gap (distance between opposing epithelial tongues) in aged wounds at 1 and 3 dpw (Fig. 1F).

**Figure 1.**
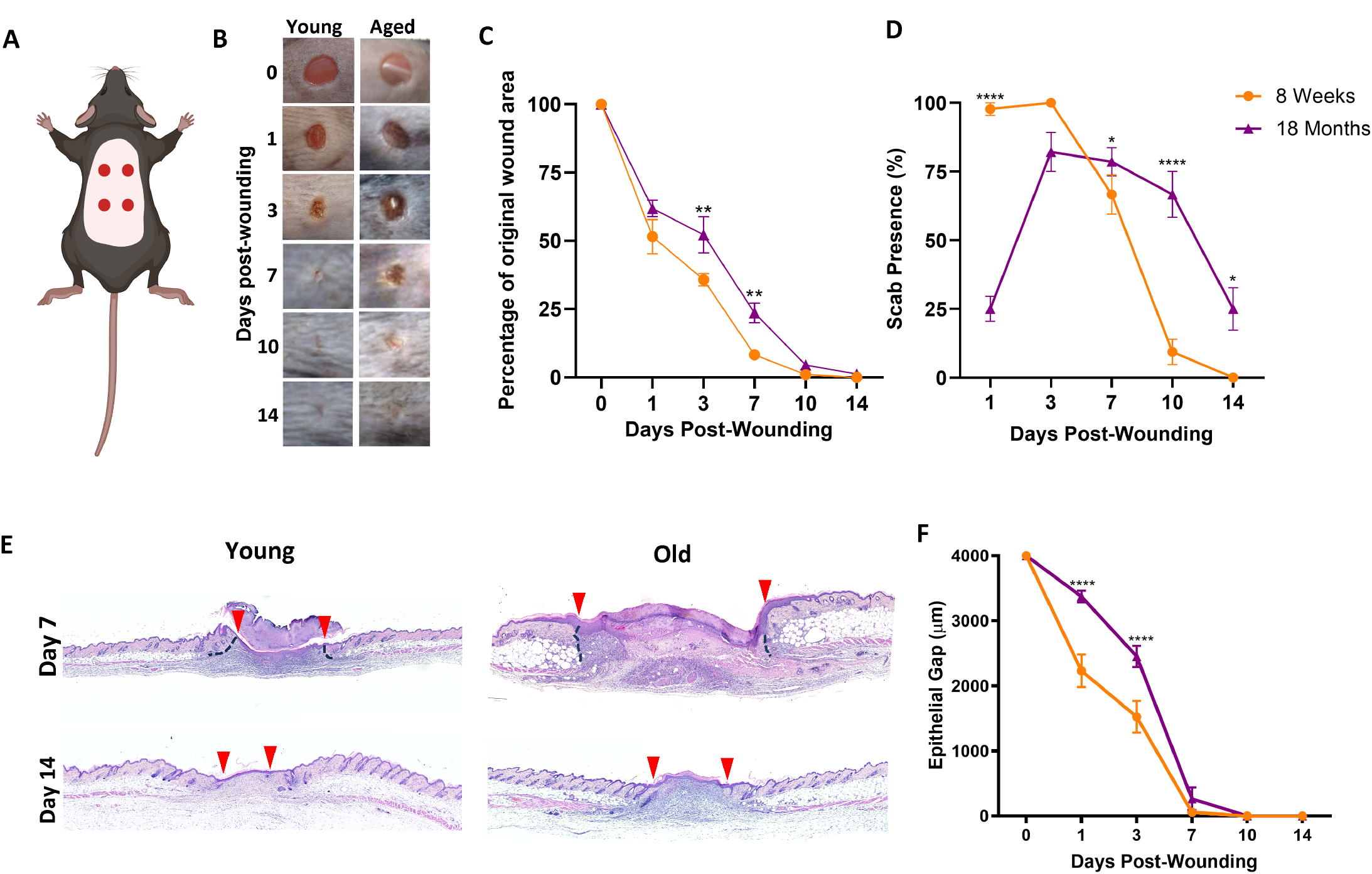
Aged skin exhibits delays throughout wound healing. **A** Schematic showing the placement of tour 4 mm excisional wounds made to the dorsal skin of young (8 week) and old (18 month) C57BI6/JCrl mice. **B** Representative macroscopic photos of young and age-impaired skin repair 0-14 days post wounding (dpw). **C** Quantification of wound areas in young and aged mice 0-14 dpw, **D** Quantification of scab presence 1-14 DPI. **E** Representative H&E-stained wound mid-sections at 7 and 14 days post-wounding in young and old mice. Red arrows indicate wound edges. Black dotted lines indicate margin of granulation tissue. **F** Quantification of epithelial gap (distance between epithelial tongues) 0-14 DPI. 0 μm is given where re-epithelialisation is complete. 7-10 biological replicates per group from 2 independent experiments. *, p<0.05; **, p<0.01; ****, p<0.0001 relative to 8 week old animals, by 2-way ANOVA with Bonferroni’s post-hoc test.

Collectively, these data confirm that advanced age is associated with delayed progression through key stages of cutaneous repair, including wound contraction and re-epithelialisation. Because this impairment reflects delayed, rather than absent, repair, we next asked whether inflammatory resolution within the wound bed was similarly prolonged.

### Neutrophils persist in age-impaired healing and display altered spatial distribution

Timely resolution of inflammation is essential for effective wound repair and depends on the coordinated clearance of myeloid cells from the wound bed. We therefore examined the temporal dynamics and spatial organisation of neutrophils and macrophages during the resolution phase of wound healing in young and aged mice. At 7 days post-wounding (dpw), both young and aged wounds contained a substantial neutrophil infiltrate; however, neutrophil density was already significantly higher in aged wounds (1.9-fold increase relative to young; Fig. 2A–B). This divergence became more pronounced at 10 dpw, with aged wounds containing 4.3-fold more neutrophils than young wounds (123 vs 29 cells/mm^2^; Fig. 2A–B). In young mice, neutrophil numbers declined significantly between 7 and 10 dpw (76 to 29 cells/mm^2^), consistent with progression toward inflammatory resolution. In contrast, no comparable reduction was observed in aged wounds, and although neutrophil numbers decreased by 14 dpw in both groups, they remained significantly elevated in aged animals.

**Figure 2.**
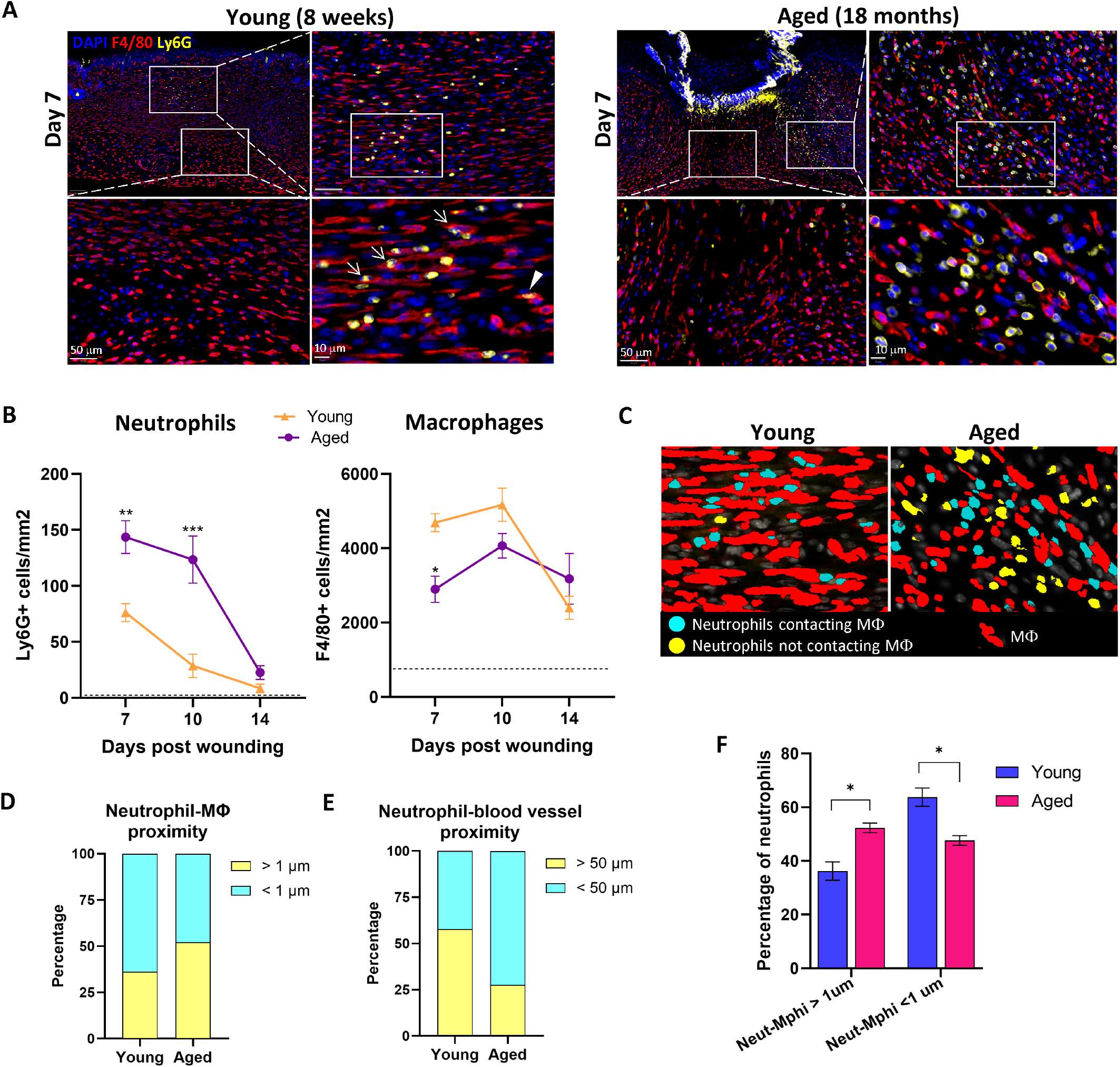
Neutrophils persist in age-impaired healing and display altered spatial distribution. **A** Representative multiplexed immunofluorescence images showing neutrophil (Ly6G; yellow) interactions with macrophages (F4/80; red) with DAPI nuclear stain in blue 7 days. **B** Quantification of wound bed neutrophils, macrophages and dendritic cells normalised to mm^2^. Dotted line indicates number of neutrophils and macrophages present in unwounded skin. *, p<0.05; **, p<0.01; ***, p<0.001 relative to Young (8 week old) wounds, by 2 way ANOVA with Sidaks post-hoc test. **C** Image analysis in Visiopharm depicting the interaction between neutrophils and macrophages. Neutrophils in contact with macrophages (cyan), neutrophils not contacting macrophage (yellow), and macrophages (red) masks. Macrophage mask was expanded by 1 μm from cell boundary and neutrophils within 1 μm of macrophage considered in contact. **D, E** Stacked bar graphs illustrating percentage **(D)** neutrophils within 1 μm boundary of macrophages (cyan) and neutrophils outside 1 μm boundary of macrophages (yellow) **(E)** neutrophils within the perivascular niche defined as 50 μm from blood vessel boundary (cyan) and neutrophils outside the perivascular niche (yellow) in young and aged mice. **F** Percentage of neutrophils not in close proximity and in close proximity with macrophages (<1 μm). *, p<0.05 by 2-way ANOVA.

Macrophage abundance displayed a distinct age-dependent pattern. At 7 dpw, macrophages were more abundant in young wounds compared with aged wounds, followed by a significant reduction by 14 dpw; this temporal pattern was not observed in aged animals (Fig. 2A–B).

Spatial analysis revealed marked differences in neutrophil–macrophage organisation between groups. In young wounds, the majority of neutrophils were found in direct contact with macrophages, consistent with ongoing clearance processes (Fig. 2C–D, F). In contrast, in aged wounds neutrophils were predominantly localised around blood vessels at the wound margins, with reduced proximity to macrophages (Fig. 2C, E–F), suggestive of continued recruitment and/or impaired dispersal within the wound bed.

### Persistent inflammatory and chemokine gene expression characterises age-impaired wound repair

To investigate molecular programmes associated with persistent neutrophil accumulation in aged wounds, we performed focused transcriptional profiling of wounds using the NanoString Myeloid Gene Expression Panel at 1, 3 and 7 days post-injury (dpw). Differential expression analysis revealed substantial overlap in wound-induced transcriptional responses between young and aged animals at all time points, alongside a smaller but consistent set of age-specific differentially expressed genes (DEGs; Fig. 3A).

**Figure 3.**
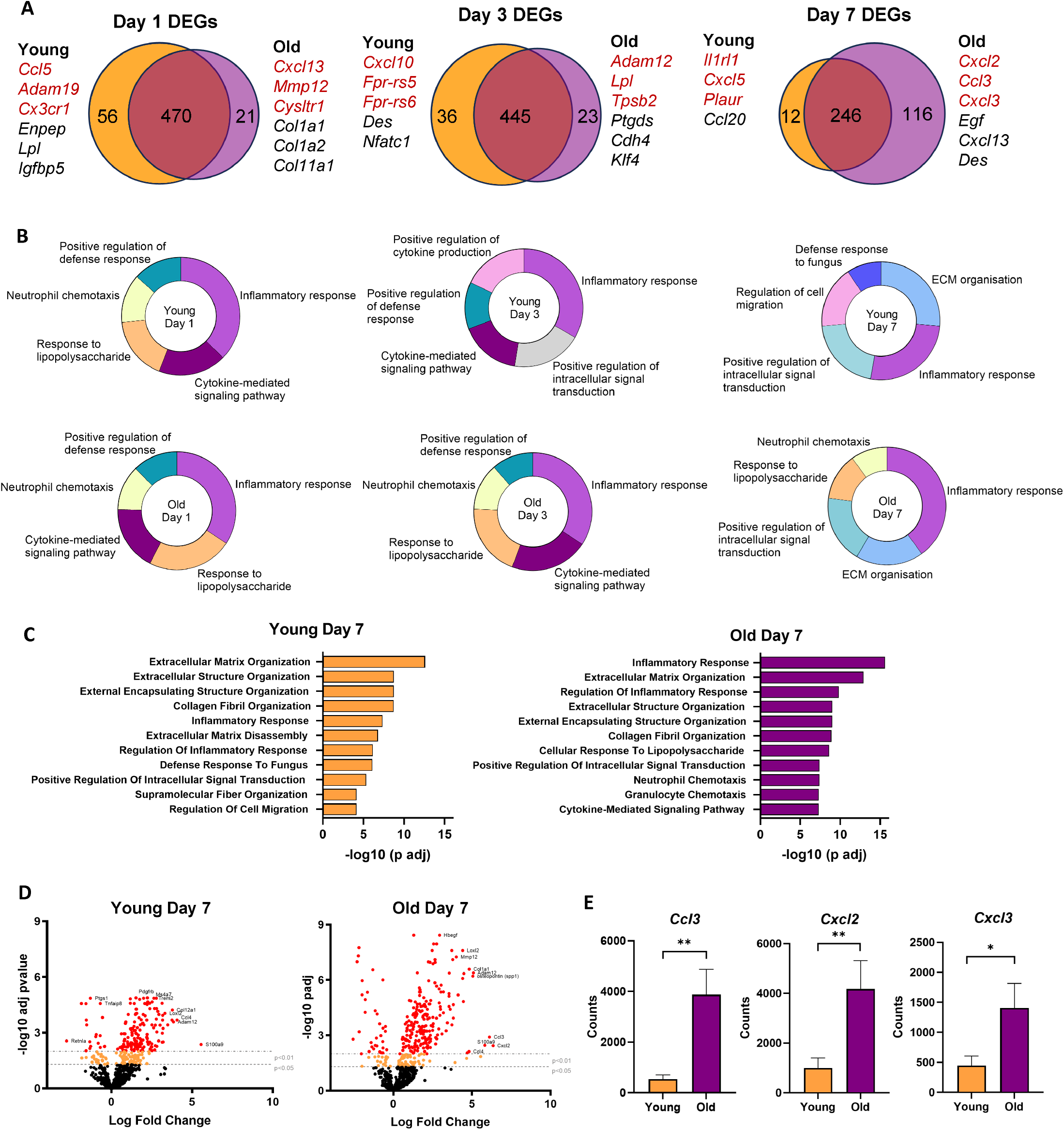
Persistent inflammatory and chemokine gene expression characterises age-impaired wound repair. **A**. Venn diagrams illustrating shared and unique DEGs between young and aged animals at days 1, 3 and 7 post-wounding. Top 3 unique upregulated genes (red) and top 3 unique downregulated genes (black) for each age and time point. **B** Percentage of top 50 DEGs that associated with the GO terms. GO BP analysis performed with enrichr. Top 5 terms p<0.05 and containing at least 4 genes are shown. GO terms combined where gene lists were identical. **C** GO terms associated with top 50 DEGs. **D** Volcano plots of wounds young and age-impaired wound repair at 7 dpi normalised to young and aged unwounded skin. Red and orange dots represent DEGs with significance lines shown. **E** Expression of *Ccl3, Cxcl2* and *Cxcl3* transcripts in young (n=11) and age-impaired (n= 9) skin repair 7 dpi. **, p<0.01; *, p<0.05 by Students t test.

At early time points (1 and 3 dpw), both young and aged wounds exhibited enrichment of inflammatory and cytokine-mediated signalling pathways, consistent with an acute innate immune response to injury (Fig. 3B, Supplementary Table 1). However, by 7 dpw, transcriptional programmes diverged markedly between age groups. In young wounds, DEGs were predominantly associated with extracellular matrix organisation, collagen fibril organisation and resolution-associated processes, alongside attenuation of inflammatory signalling (Fig. 3B–C). In contrast, aged wounds retained strong enrichment for inflammatory response terms, including neutrophil and granulocyte chemotaxis, cytokine-mediated signalling and responses to inflammatory stimuli (Fig. 3B–C).

Direct comparison of DEGs at 7 dpw further highlighted this divergence, with aged wounds exhibiting a greater magnitude and number of inflammation-associated transcripts relative to young wounds (Fig. 3D). Notably, several chemokines implicated in neutrophil recruitment were selectively upregulated in aged wounds, including *Ccl3, Cxcl2* and *Cxcl3* (Fig. 3A, D).

Consistent with these transcriptomic findings, quantitative analysis confirmed significantly elevated expression of *Ccl3, Cxcl2* and *Cxcl3* in aged compared with young wounds at 7 dpw (Fig. 3E). Together, these data indicate that age-impaired wound repair is associated with sustained activation of myeloid inflammatory transcriptional programmes and persistent expression of neutrophil-recruiting chemokines during a phase when inflammation is resolving in young animals.

### CCL3 blockade rescues age-impaired healing by limiting sustained neutrophil recruitment

To determine whether persistent CCL3 signalling contributes functionally to delayed wound repair in aged animals, we first assessed the cellular sources of *CCL3* and the related neutrophil chemokine *CXCL2* using publicly available single-cell RNA sequencing datasets from healthy human skin and venous leg ulcers (Supplementary Figure 1). Both chemokines were predominantly expressed by myeloid populations, with enrichment in macrophage clusters in ulcerated tissue compared with healthy skin (Fig. 4A–B), consistent with a wound-associated inflammatory source.

**Figure 4.**
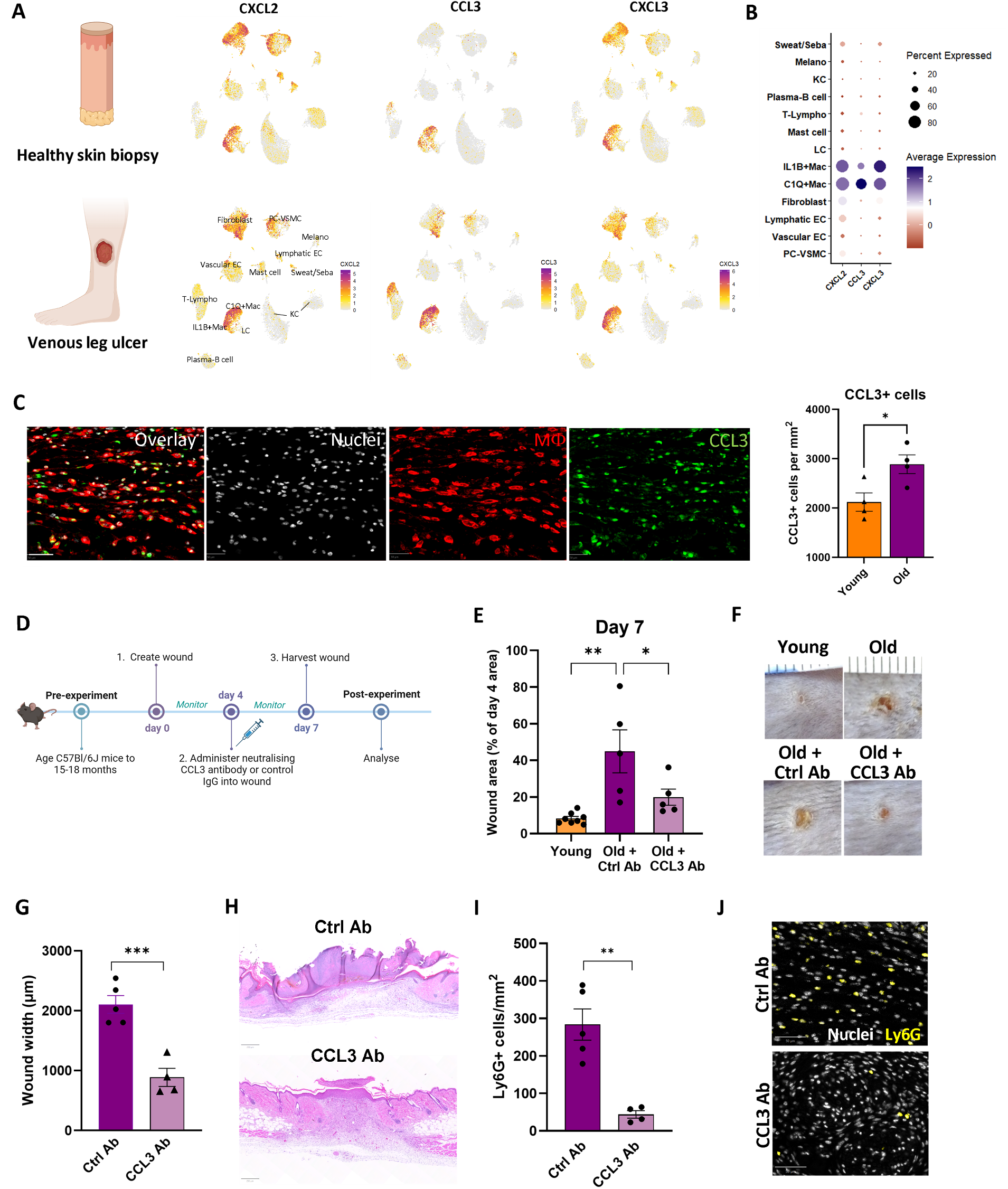
CCL3 blockade rescues age impaired healing by limiting sustained neutrophil recruitment. **A**. Feature plot depicting the transcript expression of CCL3, CXCL2 and CXCL3 across different cell population in healthy skin and venous leg ulcer (VLU) dataset. **B**. Dot plot showing expression of CCL3, CXCL2 and CXCL3 (x axis), plotted against cell types in healthy skin and VLU (y axis). Size of dots represents the fraction of cells expressing a specific marker, and color intensity indicates average expression levels; red: low, purple: high. **C** Representative multiplex immunofluorescence images illustrating the colocalization of F4/80+ macrophages (red) and CCL3+ cells (green). DAPI nuclear stain in white. Right: Quantification of CCL3+ cells normalised to mm^2^ in young (n=4) and old (n=4) mice. *, p<0.05 by Students t test. **D** Schematic of CCL3 blockade experiment. **E** Quantification of wound area in young (n=8), Old+Ctrl Ab (n= 5) and Old+CCL3 Ab (n=5) mice at 7 dpw. 1 way ANOVA with Bonferonni post-hoc test. *, p<0.05; **, p<0.01. **F** Representative macroscopic photos of day 7 wounds in young mice comparing to old mice and CCL3 Ab-treated wounds in comparison to Ctrl Ab wounds in old mice. **G** Quantification of wound width at 7 dpw in Ctrl Ab wounds (n=5) and CCL3 Ab wounds (n=4). ***, p<0.001 by Students t test. **H** Representative H&E-stained wound mid-sections at 7 dpw illustrating smaller wound area in CCL3 Ab-treated wounds. **I** Number of Ly6G+ cells/mm^2^ in Ctrl Ab and CCL3 Ab-treated wounds 7 dpw. **, p<0.01 by Students t test. **J** Representative immunofluorescence images showing reduction in wound neutrophil infiltration in CCL3 Ab-treated wounds. Keratinocytes (KC), endothelial cells (EC), fibroblasts, pericytes/vascular smooth muscle cells (PC-VSMC), myeloid subsets (C1Q+ and IL1b+ macrophages and Langerhans cells; LC), lymphoid populations (T-lymphocytes, Plasma B cells), mast cells, melanocytes (Melano) and sweat/sebaceous gland cells (Sweat/Seba). Ab, antibody; Ctrl, control.

In murine wounds, multiplex immunofluorescence confirmed increased abundance of CCL3^+^ cells in aged compared with young wounds at 7 dpw, with CCL3 expression localising predominantly to macrophages within the wound bed (Fig. 4C). Quantitative analysis revealed a significant increase in CCL3^+^ cells per mm^2^ in aged wounds relative to young controls (Fig. 4C).

To directly test whether sustained CCL3 signalling contributes to impaired healing in aged mice, we administered a neutralising anti-CCL3 antibody during the inflammatory phase of wound repair (Fig. 4D). Blockade of CCL3 significantly reduced wound area at 7 dpw in aged mice compared with age-matched control antibody–treated animals, partially restoring wound closure toward levels observed in young mice (Fig. 4E–F). Consistent with improved macroscopic healing, CCL3 blockade resulted in a marked reduction in wound width and improved epithelial architecture at 7 dpw, as assessed by histological analysis (Fig. 4G–H).

Given the established role of CCL3 in neutrophil recruitment, we next assessed neutrophil abundance following CCL3 neutralisation. Aged wounds treated with anti-CCL3 antibody exhibited a significant reduction in Ly6G^+^ neutrophils within the wound bed compared with control-treated aged wounds (Fig. 4I–J). These findings indicate that sustained CCL3 signalling actively contributes to prolonged neutrophil accumulation and delayed wound repair in advanced age, and that interrupting this axis is sufficient to improve healing outcomes.

## DISCUSSION

Ageing is a dominant risk factor for delayed and non-healing cutaneous wounds, yet the mechanistic basis for this vulnerability remains incompletely resolved. Much of the existing literature has focused on intrinsic defects in keratinocytes, fibroblasts, or vascular function with age, while comparatively less attention has been paid to how ageing alters the *resolution* phase of inflammation. The present study advances this understanding by identifying sustained myeloid-derived chemokine signalling, specifically CCL3, as an active driver of prolonged neutrophil recruitment and delayed repair in aged skin.

Previous work has established that neutrophils are essential for early wound defence but become detrimental when retained beyond the acute inflammatory phase, contributing to tissue damage, protease excess and impaired re-epithelialisation. Ageing has been associated with increased neutrophil burden in wounds, but this has largely been interpreted as a passive consequence of delayed healing rather than a regulated process. Our data instead support a model in which ageing disrupts the normal transition from inflammatory to reparative programmes, resulting in continued chemokine-mediated neutrophil recruitment at a time when inflammation is resolving in young tissue.

Our findings are consistent with a recent single-cell transcriptomic analysis of young and aged murine skin wounds by Vu *et al*. (*Cell Reports*, 2022), which identified persistent neutrophil populations in aged wounds alongside sustained inflammatory macrophage programmes. That study provided important systems-level evidence that ageing disrupts immune resolution during repair. Our data extend these observations by combining temporal, spatial and functional analyses to demonstrate that neutrophil persistence in aged wounds reflects ongoing chemokine-mediated recruitment and that sustained CCL3 signalling actively contributes to delayed healing.

The perivascular localisation of neutrophils in aged wounds, together with sustained expression of neutrophil-attracting chemokines, is consistent with ongoing recruitment from the circulation rather than simple failure of clearance. This distinction has important implications, as it suggests that therapeutic strategies aimed at modulating inflammatory *signalling*—rather than broadly suppressing immune cells—may be sufficient to restore more effective healing dynamics in aged tissue.

CCL3 has previously been implicated in leukocyte recruitment in inflammatory and fibrotic contexts, including chronic inflammatory skin disease and impaired healing in metabolic disorders. However, its role in age-associated wound repair has not been functionally tested. By demonstrating that transient CCL3 blockade during the inflammatory phase improves healing outcomes in aged mice, our study identifies CCL3 as a nodal regulator linking macrophage inflammatory programmes to sustained neutrophil accumulation in ageing skin.

Notably, CCL3 neutralisation did not ablate inflammation, but rather attenuated its persistence, supporting the concept that ageing skews inflammatory *duration* rather than magnitude.

The identification of macrophages as a major source of CCL3 in aged wounds further aligns with emerging evidence that ageing alters macrophage phenotype, persistence and inflammatory tone. Age-associated changes in macrophage metabolism, efferocytotic capacity and cytokine production have all been described and may converge to promote a self-sustaining inflammatory niche. Whether macrophage-derived CCL3 directly interferes with neutrophil clearance, promotes repeated recruitment, or both, remains an important question for future work.

Several limitations should be considered. Our transcriptional analyses were performed on whole wound tissue using a myeloid-focussed panel rather than purified populations, limiting cell-type resolution. In addition, while spatial proximity analyses suggest altered neutrophil– macrophage interactions, direct assessment of efferocytosis in aged wounds will be required to fully define the mechanisms underlying neutrophil persistence. Finally, although CCL3 expression localised predominantly to macrophages within aged wounds, antibody-mediated blockade cannot definitively assign the relative contribution of individual cellular sources to the observed repair phenotype.

Despite these limitations, this work has translational relevance. Chronic wounds in older individuals, including diabetic foot ulcers and venous leg ulcers, are characterised by prolonged inflammation and excessive neutrophil infiltration. The demonstration that targeting a single chemokine axis can rebalance inflammatory resolution without preventing wound closure highlights the potential for temporally targeted immunomodulatory therapies in ageing-associated wound pathology.

In conclusion, our findings support a model in which ageing sustains inflammatory chemokine signalling within the wound microenvironment, prolonging neutrophil recruitment and delaying repair. By identifying CCL3 as a modifiable regulator of this process, this study provides new mechanistic insight into age-impaired wound healing and identifies inflammatory resolution as a promising therapeutic entry point.

## MATERIALS AND METHODS

Key reagents and resources are detailed in Supplementary Table 2.

### Animals

All experiments were conducted with approval from the University of Edinburgh Local Ethical Review Committee and in accordance with the UK Home Office regulations (Guidance on the Operation of Animals, Scientific Procedures Act, 1986) under PPL PD3147DAB. C57Bl/6JCrl were bred in-house (LFR at BioQuarter site), maintained in conventional cages on a 12:12 light:dark cycle with *ad libitum* access to standard chow and water under an SPF environment. Animals were housed 3-5 per cage in a temperature (22-24C) and humidity-controlled room. All animals were housed in the same room throughout their lifespan. Environmental enrichment was provided in the form of dome homes, a tunnel and wooden chew sticks. Health checks were performed on all animals prior to wounding, including baseline weight measurements, malocclusion and skin injuries due to fighting. Animals with malocclusion were excluded from the study and those with small skin injuries due to historic fighting were included providing wounds could be made without involvement of the previous injury area. Only healthy animals that were not involved in previous procedures were used for experiments. The acute wound model is categorized as a moderate severity protocol.

### Aging mouse colony

Once animals reached 12 months of age, several parameters of health were monitored on a weekly basis to determine animal welfare. Parameters were weight (as compared to baseline 12 month weight), body condition scores (as determined by visual and physical examination of each animal in accordance with the University of Edinburgh Veterinary Scientific Service guidelines) and behaviour. Weight loss of 10% and/or an increase in body condition score or evident lack of expected behaviour (interaction, movement) resulted in the animal being assessed by a veterinary surgeon.

### Cutaneous wounding

Male and female mice at either 8 weeks old (Young) or 18 months old (Old) were randomly assigned a group and anaesthetized with isoflurane (Zoetis, Leatherhead, UK) by inhalation. Buprenorphine analgesia (0.05 mg/kg, s.c, Vetergesic, Amsterdam) was provided immediately prior to wounding and dorsal hair was removed using a Wahl trimmer. Four full-thickness excisional wounds were made to the shaved dorsal skin using sterile, single-use 4 mm punch biopsy tools (Selles Medical, Hull, UK). Wounds were photographed with a Sony DSC-WX350 and a ruler immediately after wounding and also on days 1, 3, 7, 10 and 14 post-wounding. Wound areas were calculated using ImageJ and normalised to day 0 areas. Mice were housed with their previous cage mates in a 28C warm box (Scanbur, Denmark) overnight following wounding, with paper towels used as bedding to avoid sawdust entering the open wounds. Dome home entrances were enlarged to minimise contact with dorsal wounds. The following day, animals were moved into clean conventional cages at 22-24C. Wounds were made in telogen-phase skin. Any wounds in which the surrounding hair follicles transitioned to catagen or anagen during the experimental period were excluded from analysis due to the established influence of hair cycle stage on wound healing (14).

For CCL3 blocking experiments, mice were anaesthetized with isoflurane by inhalation 4 days post-wounding and 5 μl anti-CCL3 neutralizing antibody (ThermoFisher Scientific, PA5-46951; 1 μg/ml) or IgG control (ThermoFisher Scientific; 1 μg/ml) were injected into each wound bed using a 30 G needle (Becton Dickinson). Animals were allowed to recover from anaesthesia in a 28C warm box before being returned to room temperature (22-24C).

### Wound harvesting and tissue processing

Acute wounds were harvested on days 1, 3, 7, 10 and 14 post-wounding, fixed in 4% PFA (overnight at 4C on a rocker, Sigma Aldrich), washed 3 × 5 min in PBS, transferred to 70% ethanol and then embedded in paraffin. Day 1 and 3 wounds were cut in half using a scalpel and sections (10 μm) taken from the middle of the wound. Day 7, 10 and 14 wounds were sectioned through the wound beyond the midpoint and wound centres identified by staining with Haematoxylin (Gills No.3, Sigma Aldrich) and Eosin (Sigma Aldrich).

### Haematoxylin and Eosin staining

Formalin-fixed, paraffin-embedded tissue sections were deparaffinised, rehydrated, and stained with haematoxylin and eosin (H&E) using standard protocols. Sections were dehydrated, cleared, and mounted with DPX mounting medium (Sigma Aldrich) prior to imaging.

### Opal Multiplex Immunofluorescence and PhenoImager HT

Multiplex immunofluorescence was performed on formalin-fixed, paraffin-embedded skin sections using the Opal™ tyramide signal amplification (TSA) system (Akoya Biosciences) to detect F4/80, Ly6G, CCL3 and CD31. Sections were deparaffinised, rehydrated and subjected to heat-mediated antigen retrieval, followed by blocking and sequential staining cycles according to the manufacturer’s instructions (Akoya Biosciences). Primary antibodies against F4/80, Ly6G, CCL3 and CD31 (suppliers, catalogue numbers and dilutions listed in Supplementary Table 2) were applied at optimised concentrations and detected using HRP-conjugated secondary reagents (Akoya Biosciences) with Opal fluorophores 520, 570, 690 and 780 (Akoya Biosciences). A TSA-DIG amplification step (Akoya Biosciences) was incorporated in the final staining cycle. Nuclei were counterstained with Spectral DAPI (Akoya Biosciences), and sections were mounted using ProLong™ Diamond Antifade Mountant (ThermoFisher Scientific, UK).

Whole-slide fluorescence imaging was performed using the PhenoImager HT system (Akoya Biosciences) at 40× magnification. Regions of interest were selected using Phenochart software (Akoya Biosciences), followed by spectral unmixing in inForm software (Akoya Biosciences) and image stitching and downstream analysis using Visiopharm software (Visiopharm, Hørsholm, Denmark).

### Image analysis in Visiopharm

Visiopharm (version 2023.09.4.15595 x64) was utilised to analyse the images from multiplexed immunofluorescence staining. Each tissue sample was segmented into region of interest (ROI), and only the cells within these ROIs were quantified. The ROI area was measured using the area measurement function and subsequent cell counts normalised to mm^2^. For cell detection, a nuclei detection application utilising a deep learning approach was employed to identify nuclei. A threshold classification method was applied to identify the minimum intensity of each biomarker, specifically neutrophils, macrophages and CCL3+ cells. To assess neutrophil-macrophage proximity, post-processing steps were implemented to refine the morphological boundaries of individual cells. Each macrophage boundary was expanded by 1 μm to identify neutrophils in close proximity. Neutrophils within this 1 μm boundary were labelled as Neut-Mphis <1 μm, indicating neutrophils in contact with macrophages. Those beyond this boundary were labelled Neut-Mphi >1 μm. Finally, the labelled cells were quantified using the applications count function in the output variables. Neutrophil-blood vessel proximity was defined using a threshold classification method based on the CD31 biomarker and a dilation function to expand the vessel area by 50 μm.

### RNA extraction and NanoString gene expression profiling

RNA was extracted from whole wound tissue using the RNeasy Mini Kit (Qiagen) according to the manufacturer’s instructions. Gene expression profiling was performed using the NanoString nCounter Myeloid Innate Immunity Panel which includes 770 genes. NanoString gene expression data were analysed using the ROSALIND platform (https://rosalind.bio/; ROSALIND Inc., San Diego, CA, USA). Quality control metrics included read distribution, violin plots, identity heatmaps and multidimensional scaling. Data were normalised using the nCounter Advanced Analysis workflow, with counts divided by the geometric mean of selected normaliser probes. Housekeeping genes were selected using the geNorm algorithm implemented in the NormqPCR R package. Differential expression and statistical significance were calculated using the nCounter Advanced Analysis 2.0 methodology, with p-values adjusted using the Benjamini–Hochberg false discovery rate. GO BP analysis was performed using enrichr (https://maayanlab.cloud/Enrichr/) (15). This dataset will be deposited on GEO on acceptance of the manuscript.

### Data analysis of GSE265972

The investigation utilized pre-processed and filtered scRNA-seq datasets originally published by Liu Z *et al*. (16), importing them into R environment (version 4.3.1) for independent analysis. Data normalization employed the SCTransform methodology available in Seurat Bioconductor (v5.1), applying regularized negative binomial modelling to address the characteristic sparsity of single-cell data. Following normalization and merging of expression profiles, both unsupervised and supervised analytical approaches were implemented through diverse R and Bioconductor computational tools. Stringent quality control parameters excluded cells exhibiting mitochondrial gene representation exceeding 10%, cells with gene expression below 200, and genes detected in fewer than 3 cells. Dimensionality reduction through principal component analysis identified key variance-capturing components, which subsequently facilitated Uniform Manifold Approximation and Projection (UMAP) visualization of cellular relationships. The shared nearest neighbour (SNN) approach enabled identification of transcriptionally similar cell clusters, which underwent marker detection procedures. Cell type annotation relied on expression patterns of established marker transcripts, consistent with classifications reported in the original publication (16).

### Diagrams

BioRender was used to create the schematics in Figures 1 and 4.

### Statistical analysis

Student’s *t*-test, one-way and two-way ANOVA with Bonferroni’s multiple comparison test were performed using GraphPad Prism 10.2.0 software and detailed in the respective Figure legends.

## Acknowledgments

We acknowledge the support of the IRR SURF facilities for histological services and BVS (Bioresearch and Veterinary Services), especially Dr Nacho Vinuela-Fernandez for assistance with animal studies. We thank Dr Alison Munro of the Institute of Genetics and Cancer (IGC) for running nanostring assays. Thanks also to Prof Stuart Forbes, Prof David Dockrell and Prof Julia Dorin for mentorship.

## CRediT Contributions

HR: Data Curation, Formal Analysis, Investigation, Resources, Visualisation, Writing. YN: Data Curation, Formal Analysis, Investigation, Visualisation, Writing. VS and AP: Data Curation. SRW: Supervision. JLC: Conceptualisation, Data Curation, Formal Analysis, Funding Acquisition, Investigation, Supervision, Validation, Visualisation, Writing.

