## Supplemental Table 1 for "Persistent CCL3 expression sustains neutrophil recruitment and contributes to delayed wound healing in advanced age"

| Day 1 Young | | |
|---|---|---|
| **GO Term Biological Process** | p adj | -log10 padj |
| Inflammatory Response (GO:0006954) | 1.24E-30 | 29.91 |
| Regulation Of Inflammatory Response (GO:0050727) | 2.33E-13 | 12.63 |
| Neutrophil Chemotaxis (GO:0030593) | 3.02E-13 | 12.52 |
| Cytokine-Mediated Signaling Pathway (GO:0019221) | 3.02E-13 | 12.52 |
| Granulocyte Chemotaxis (GO:0071621) | 3.35E-13 | 12.47 |
| Neutrophil Migration (GO:1990266) | 4.90E-13 | 12.31 |
| Positive Regulation Of Inflammatory Response (GO:0050729) | 9.51E-12 | 11.02 |
| Cellular Response To Lipopolysaccharide (GO:0071222) | 4.46E-11 | 10.35 |
| Positive Regulation Of Defense Response (GO:0031349) | 4.46E-11 | 10.35 |
| Positive Regulation Of Cytokine Production (GO:0001819) | 6.11E-11 | 10.21 |
| Positive Regulation Of Response To External Stimulus (GO:0032103) | 3.27E-10 | 9.49 |

| Day 1 Old | | |
|---|---|---|
| **GO Term Biological Process** | p adj | -log10 padj |
| Inflammatory Response (GO:0006954) | 4.89E-25 | 24.31 |
| Cellular Response To Lipopolysaccharide (GO:0071222) | 3.79E-12 | 11.42 |
| Cytokine-Mediated Signaling Pathway (GO:0019221) | 1.12E-11 | 10.95 |
| Neutrophil Chemotaxis (GO:0030593) | 1.94E-11 | 10.71 |
| Granulocyte Chemotaxis (GO:0071621) | 2.03E-11 | 10.69 |
| Response To Lipopolysaccharide (GO:0032496) | 2.03E-11 | 10.69 |
| Neutrophil Migration (GO:1990266) | 2.72E-11 | 10.57 |
| Positive Regulation Of Inflammatory Response (GO:0050729) | 3.85E-10 | 9.41 |
| Cellular Response To Molecule Of Bacterial Origin (GO:0071219) | 1.00E-09 | 9.00 |
| Regulation Of Inflammatory Response (GO:0050727) | 1.10E-09 | 8.96 |
| Positive Regulation Of Defense Response (GO:0031349) | 1.39E-09 | 8.86 |

| Day 3 Young | | |
|---|---|---|
| **GO Term Biological Process** | p adj | -log10 padj |
| Inflammatory Response (GO:0006954) | 7.64E-27 | 26.12 |
| Regulation Of Inflammatory Response (GO:0050727) | 7.57E-12 | 11.12 |
| Positive Regulation Of Cytokine Production (GO:0001819) | 8.27E-12 | 11.08 |
| Cytokine-Mediated Signaling Pathway (GO:0019221) | 9.15E-12 | 11.04 |
| Positive Regulation Of Inflammatory Response (GO:0050729) | 1.31E-11 | 10.88 |
| Positive Regulation Of Defense Response (GO:0031349) | 6.60E-11 | 10.18 |
| Positive Regulation Of Intracellular Signal Transduction (GO:1902533) | 1.80E-10 | 9.74 |
| Positive Regulation Of Response To External Stimulus (GO:0032103) | 4.31E-10 | 9.37 |
| Regulation Of Neuroinflammatory Response (GO:0150077) | 4.31E-10 | 9.37 |
| Neutrophil Chemotaxis (GO:0030593) | 5.83E-10 | 9.23 |
| Granulocyte Chemotaxis (GO:0071621) | 7.50E-10 | 9.12 |

| Day 3 Old | | |
|---|---|---|
| **GO Term Biological Process** | p adj | -log10 padj |
| Inflammatory Response (GO:0006954) | 4.79E-25 | 24.32 |
| Cytokine-Mediated Signaling Pathway (GO:0019221) | 1.58E-14 | 13.80 |
| Cellular Response To Lipopolysaccharide (GO:0071222) | 2.47E-12 | 11.61 |
| Neutrophil Chemotaxis (GO:0030593) | 1.90E-11 | 10.72 |
| Granulocyte Chemotaxis (GO:0071621) | 1.98E-11 | 10.70 |
| Response To Lipopolysaccharide (GO:0032496) | 1.98E-11 | 10.70 |
| Neutrophil Migration (GO:1990266) | 2.66E-11 | 10.58 |
| Cellular Response To Molecule Of Bacterial Origin (GO:0071219) | 1.11E-09 | 8.95 |
| Cellular Response To Lipid (GO:0071396) | 1.72E-08 | 7.76 |
| Regulation Of Inflammatory Response (GO:0050727) | 2.55E-08 | 7.59 |
| Positive Regulation Of Defense Response (GO:0031349) | 5.02E-08 | 7.30 |

| Day 7 Young | | |
|---|---|---|
| **GO Term Biological Process** | p adj | -log10 padj |
| Extracellular Matrix Organization (GO:0030198) | 2.60E-13 | 12.59 |
| Extracellular Structure Organization (GO:0043062) | 1.83E-09 | 8.74 |
| External Encapsulating Structure Organization (GO:0045229) | 1.83E-09 | 8.74 |
| Collagen Fibril Organization (GO:0030199) | 1.99E-09 | 8.70 |
| Inflammatory Response (GO:0006954) | 4.71E-08 | 7.33 |
| Extracellular Matrix Disassembly (GO:0022617) | 1.57E-07 | 6.80 |
| Regulation Of Inflammatory Response (GO:0050727) | 7.27E-07 | 6.14 |
| Defense Response To Fungus (GO:0050832) | 8.04E-07 | 6.09 |
| Positive Regulation Of Intracellular Signal Transduction (GO:1902533) | 4.36E-06 | 5.36 |
| Supramolecular Fiber Organization (GO:0097435) | 6.90E-05 | 4.16 |
| Regulation Of Cell Migration (GO:0030334) | 6.90E-05 | 4.16 |

| Day 7 Old | | |
|---|---|---|
| **GO Term Biological Process** | p adj | -log10 padj |
| Inflammatory Response (GO:0006954) | 2.38E-16 | 15.6 |
| Extracellular Matrix Organization (GO:0030198) | 1.32E-13 | 12.9 |
| Regulation Of Inflammatory Response (GO:0050727) | 1.49E-10 | 9.8 |
| Extracellular Structure Organization (GO:0043062) | 1.12E-09 | 9.0 |
| External Encapsulating Structure Organization (GO:0045229) | 1.12E-09 | 9.0 |
| Collagen Fibril Organization (GO:0030199) | 1.35E-09 | 8.9 |
| Cellular Response To Lipopolysaccharide (GO:0071222) | 2.38E-09 | 8.6 |
| Positive Regulation Of Intracellular Signal Transduction (GO:1902533) | 3.61E-08 | 7.4 |
| Neutrophil Chemotaxis (GO:0030593) | 3.80E-08 | 7.4 |
| Granulocyte Chemotaxis (GO:0071621) | 4.62E-08 | 7.3 |
| Cytokine-Mediated Signaling Pathway (GO:0019221) | 5.00E-08 | 7.3 |

**Supplementary Table 1 – Gene Ontology Biological Process enrichment analysis in young and aged wounds across time**

Gene Ontology (GO) Biology Process enrichment analysis of the top 50 differentially expressed genes identified from the NanoString myeloid panel in young (n=10) and aged (n=10) wounds at day 1, day 3 and day 7 post-injury. For each age and time point, the top enriched GO terms are shown. Adjusted p values (Benjamini-Hochberg correction) and corresponding –log10(adjusted p value) are indicated.
