## Supplemental Table 2 for "Persistent CCL3 expression sustains neutrophil recruitment and contributes to delayed wound healing in advanced age"

### Supplementary Table 2 – Key Reagent and Resource Details

| REAGENT or RESOURCE | SOURCE | IDENTIFIER |
|---|---|---|
| Antibodies | | |
| Anti-mouse CCL3 antibody | ThermoFisher Scientific | PA5-46951 |
| Anti-mouse CD31 (PECAM-1) antibody | Cell Signaling Technology | 77699 |
| Anti-mouse CXCL2 antibody | ThermoFisher Scientific | 701126 |
| Anti-mouse F4/80 antibody | Biolegend | 123102 |
| Anti-mouse Ly6G antibody | Biolegend | 127602 |
| Chemicals | | |
| Bovine Serum Albumin | Sigma-Aldrich | A2153-100G |
| Buprenorphine (Vetergesic) | Ceva Animal Health | Lot no: 2309272AB |
| Eosin Y Solution, Alcoholic, with Phloxine | Sigma-Aldrich | HT110332-1L |
| Ethanol | Fisher chemicals | 10437341 |
| Gelatin | Sigma-Aldrich | G9382 CAS: 9000-70-8 |
| Hematoxylin Solution, Gill No. 3 | Sigma-Aldrich | GHS332-1L |
| Hydrochloric acid | Sigma-Aldrich | H1758 |
| Isoflurane | Piramal Critical Care | 2800025 |
| Paraformaldehyde | Sigma-Aldrich | 441244-1KG |
| Pluronic F-127 | Sigma-Aldrich | P2443-250G |
| Tris Buffered Saline | Sigma-Aldrich | 93283-100ML |
| Triton X-100 | Sigma-Aldrich | T8787-50ML |
| Tween-20 | Sigma-Aldrich | P1379-500ML |
| Xylene | Fisher chemicals | 10385910 |
| Immunohistochemistry reagents | | |
| Antigen Unmasking Solution, Citrate-Based | VectorLabs | H-3300-250 |
| Antigen Unmasking Solution, Tris-Based | VectorLabs | H-3301-250 |
| AR6 (Citrate) | Akoya Biosciences | AR600250ML |
| DPX | Fisher scientific | D/5319/05 |
| ImmEdge® Hydrophobic Barrier PAP Pen | VectorLabs | H-4000 |
| ImmPACT® DAB Substrate Kit, Peroxidase (HRP) | VectorLabs | SK-4105 |
| ImmPRESS® HRP Goat Anti-Rat IgG Polymer Detection Kit | VectorLabs | MP-7404 |
| ImmPRESS® HRP Horse Anti-Mouse IgG Polymer Detection Kit | VectorLabs | MP-7402 |
| ImmPRESS® HRP Horse Anti-Rabbit IgG Polymer Detection Kit | VectorLabs | MP-7401 |
| Normal Goat Serum | VectorLabs | S-1012 |
| Opal Polaris 7 Color IHC Detection Kits | Akoya Biosciences | OP-000003 |
| Prolong Diamond Mounting Media | Invitrogen | P36970 |
| Spectral DAPI | Akoya Biosciences | SKU FP1490 |
| Deposited data | | |
| Raw data files for single-cell RNA-seq | Liu, Z. *et al.*, 2025 | GEO: GSE265972 |
| Experimental models: Organisms/strains | | |
| C57Bl/6JCrl | Charles River | 632 |
| Software and algorithms | | |
| GraphPad Prism v10.2.0 | GraphPad Software | www.graphpad.com |
| inForm v3.0 | Akoya Biosciences | https://www.akoyabio.com/support/software/ |
| Phenochart v2.2.0 | Akoya Biosciences | https://www.akoyabio.com/support/software/ |
| Qupath v0.5.1 | Bankhead, P. et al. | https://qupath.github.io/ |
| R v4.3.1 | The R Project for Statistical Computing | https://www.r-project.org/ |
| Seurat v5.1 | Hao *et al.*, 2023 | https://satijalab.org/seurat/ |
| Visiopharm image analysis (VIS) | Visiopharm (Hoersholm, Denmark) | https://visiopharm.com/ |
| Other | | |
| 4 mm punch biopsy | Kai Industries | BP-40F |
| Coverslips (22x 60mm) 0.13 mm thick | ThermoFisher Scientific | BBAD02200600#A |
| Disposable sterile scalpels | Swann-Morton | 503 |
| SuperFrost slides | ThermoFisher Scientific | 10149870 |
