## Supplemental Figure 1 for "Persistent CCL3 expression sustains neutrophil recruitment and contributes to delayed wound healing in advanced age"

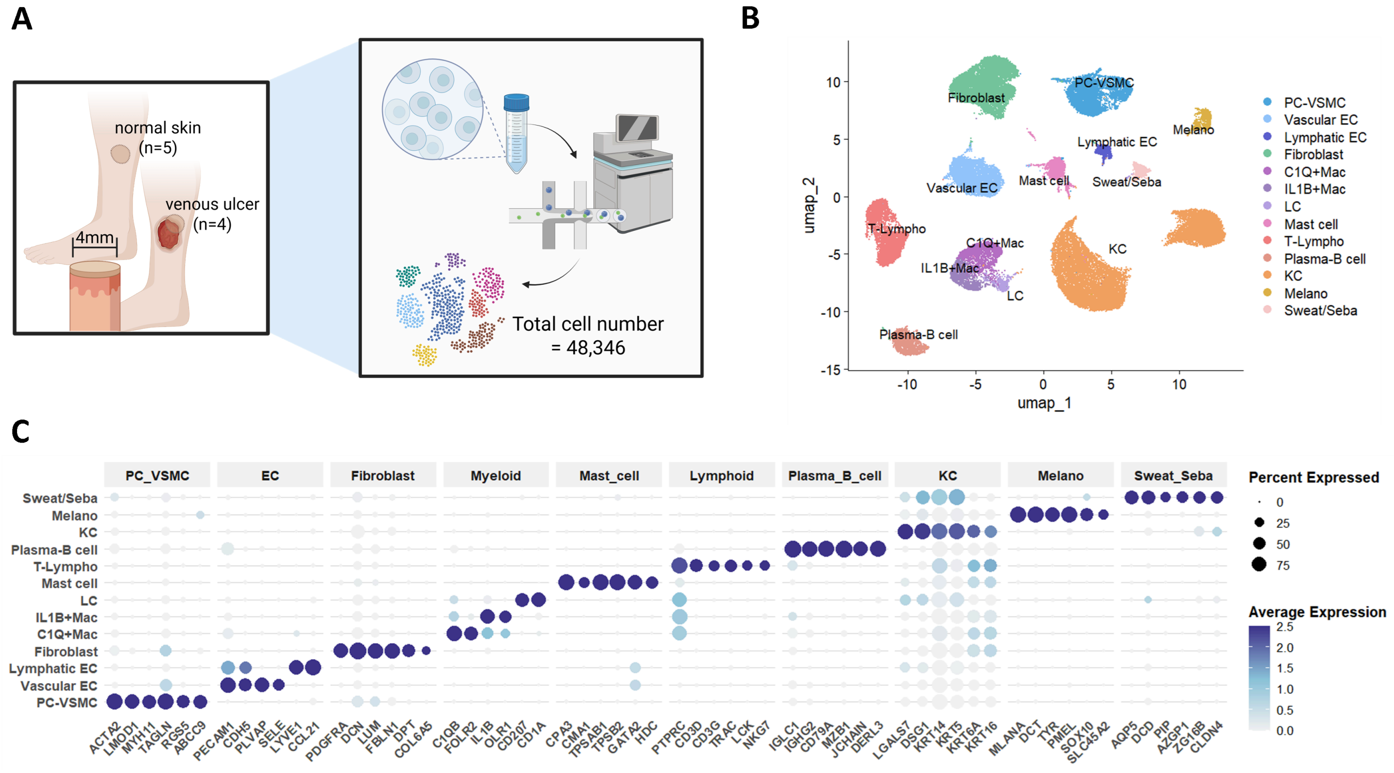


**Supplementary Figure 1 – ScRNAseq dataset overview and cell type annotation**
**A** Study overview. scRNAseq data were obtained from GSE265972 comprising normal skin (n=5) and venous ulcer tissue (n=4) biopsies. Following quality control and filtering, a total of 48,346 cells were retained for analysis. **B** UMAP embedding of all cells following clustering and annotation, coloured by major cell populations, including keratinocytes (KC), vascular endothelial cells (Vascular EC), fibroblasts, pericytes/vascular smooth muscle cells (PC-VSMC), myeloid subsets (C1Q+ and IL1b+ macrophages, Langerhans cells), lymphoid populations (T-lymphocytes, Plasma B cells), mast cells, melanocytes (Melano) and sweat/sebaceous gland cells (Sweat/Seba). **C** Dot plot showing expression of canonical marker genes used for cell type annotation. Dot size indicates the proportion of cells expressing each gene, and colour intensity represents average expression within each cluster.
